# Post-mitotic Prox1 expression controls the final specification of cortical VIP interneuron subtypes

**DOI:** 10.1101/2020.07.22.216135

**Authors:** Tevye Jason Stachniak, Rahel Kastli, Olivia Hanley, Ali Özgür Argunsah, Theofanis Karayannis

## Abstract

Neuronal identity is controlled in multiple developmental steps by key transcription factors that determine the unique properties of a cell. During embryogenesis, the transcription factor Prox1 has been shown to regulate VIP interneuron migration, survival, and as a result, circuit integration. Here, we explore the role of Prox1 as a regulator of genetic programs that guide the final specification of VIP interneuron subtypes in early post-natal life. Using in-vitro electrophysiology we find that post-natal removal of Prox1 differentially affects the synaptic integration of VIP bipolar and multipolar subtypes.

RNA sequencing reveals that one of the downstream targets of Prox1 is the postsynaptic protein Elfn1, a constitutive regulator of presynaptic release probability. Genetic, pharmacological and electrophysiological experiments demonstrate that knocking out Prox1 reduces Elfn1 function in VIP multipolar but not in bipolar cells. Thus, in addition to the activity-dependent and contextual processes that finalize developmental trajectories, genetic programs engaged by Prox1 control the differentiation and connectivity of VIP interneuron subtypes.

## Introduction

Although cortical interneuron (IN) diversity begins at their place of birth within distinct embryonic progenitor domains (Mi et al., 2018), single cell sequencing and manipulation experiments at different developmental stages have suggested that INs undergo their final specification while integrating into the developing circuit (De Marco García et al., 2011; Mayer et al., 2018). The developmental mechanisms by which distinct types of INs acquire their mature characteristics are only beginning to be revealed (Favuzzi et al., 2019). Interestingly, post-mitotic manipulations have demonstrated a persistent requirement for key transcription factors (TF) in the final specification and maintenance of pyramidal cell fate (De La Rossa et al., 2013). Whether similar TF mechanisms exist for cortical INs remains unknown.

Cortical vasoactive intestinal peptide-expressing (VIP) INs are a diverse population (Tasic et al., 2018). Even though they make up less than 5% of all neurons, VIP INs are critically important for cortical circuit maturation and their malfunction has been implicated in neurodevelopmental disorders (Batista-Brito et al., 2017; Mossner et al., 2020). The TF Prospero-related homeobox1 (Prox1) is expressed by the majority of INs derived from the caudal ganglionic eminence (CGE). Its removal during embryonic development impairs VIP cell migration, cell survival, as well as dendritic development and afferent connectivity of the Calretinin-expressing (CR+) VIP bipolar subtype (Miyoshi et al., 2015). Importantly, Prox1 remains expressed in all VIP INs as they integrate into the developing circuit, suggesting it may have a role in organizing not only their early stages of development, but also their final specification. In addition to requiring Prox1, it has been shown that CR+ VIP bipolar cells also require proper network activity to acquire their characteristic axo-dendritic profile, as do Reelin- and Somatostatin (SST)-expressing INs (De Marco García et al., 2011, 2015; Pan et al., 2018). Although Prox1 is also expressed in the cholecystokinin-expressing (CCK+) VIP multipolar subtype, these cells do not show any positional or morphological deficits following activity manipulations (De Marco García et al., 2011). Taken together, these findings raise the possibility that during the final developmental steps of VIP IN specification, cell-autonomous and activity-dependent genetic programs work in tandem to guide the network integration of distinct VIP subtypes in a differential manner.

In this study, we assess the post-natal requirement of Prox1 for the final diversification of VIP IN subtypes. To achieve this, we conditionally remove this TF during the first post-natal week, using a *VIPCre* driver mouse line, and explore the synaptic integration of bipolar and multipolar VIP cells using functional and RNA screening methods. Surprisingly, we find that Prox1 removal impacts the two subtypes differentially. Consistent with previous findings, bipolar cells have reduced synaptic excitation upon removal of Prox1. In contrast, multipolar cells show instead an alteration in the short-term synaptic dynamics of their incoming excitation. Using transcriptomic screening and pharmacological manipulations we demonstrate that a Prox1-dependent engagement of the trans-synaptic protein Elfn1 selectively enables the synaptic facilitation observed in multipolar cells.

## Results

### 1) Postnatal Prox1 removal leads to changes in presynaptic release probability onto VIP multipolar but not bipolar cells

Cell-specific synaptic wiring properties are a prominent feature of IN cell type diversity. Previous research found that embryonic Prox1 removal leads to aberrant network integration of VIP bipolar cells (Miyoshi et al., 2015) while the effect on multipolar cells is unknown. Therefore, we first wanted to test whether postnatal loss of function of Prox1 (KO) affects the network integration of bipolar, as well as multipolar VIP cells. To postnatally (∼P3) remove Prox1, we used a VIP knock-in mouse line that drives the expression of Cre from the endogenous peptide locus (*VIPCre*), combined with a conditional *Prox1* allele. The *Prox1* coding region is flanked by *loxp* sites and recombination shifts *eGFP* into frame (*Prox1eGFP)* to label all Prox1+ VIP cells (Figure 1A). We first surveyed spontaneous excitatory postsynaptic currents (sEPSCs) in L2/3 control (*Prox1* heterozygote; GFP labeled) and Prox1 KO VIP INs in acute brain slices for alterations which would indicate connectivity changes. Indeed, we found that postnatal expression of Prox1 regulates incoming excitatory inputs onto both VIP subtypes, albeit with subtype specific differences in time course and valence (Supplementary Figure 1).

**Figure 1:**
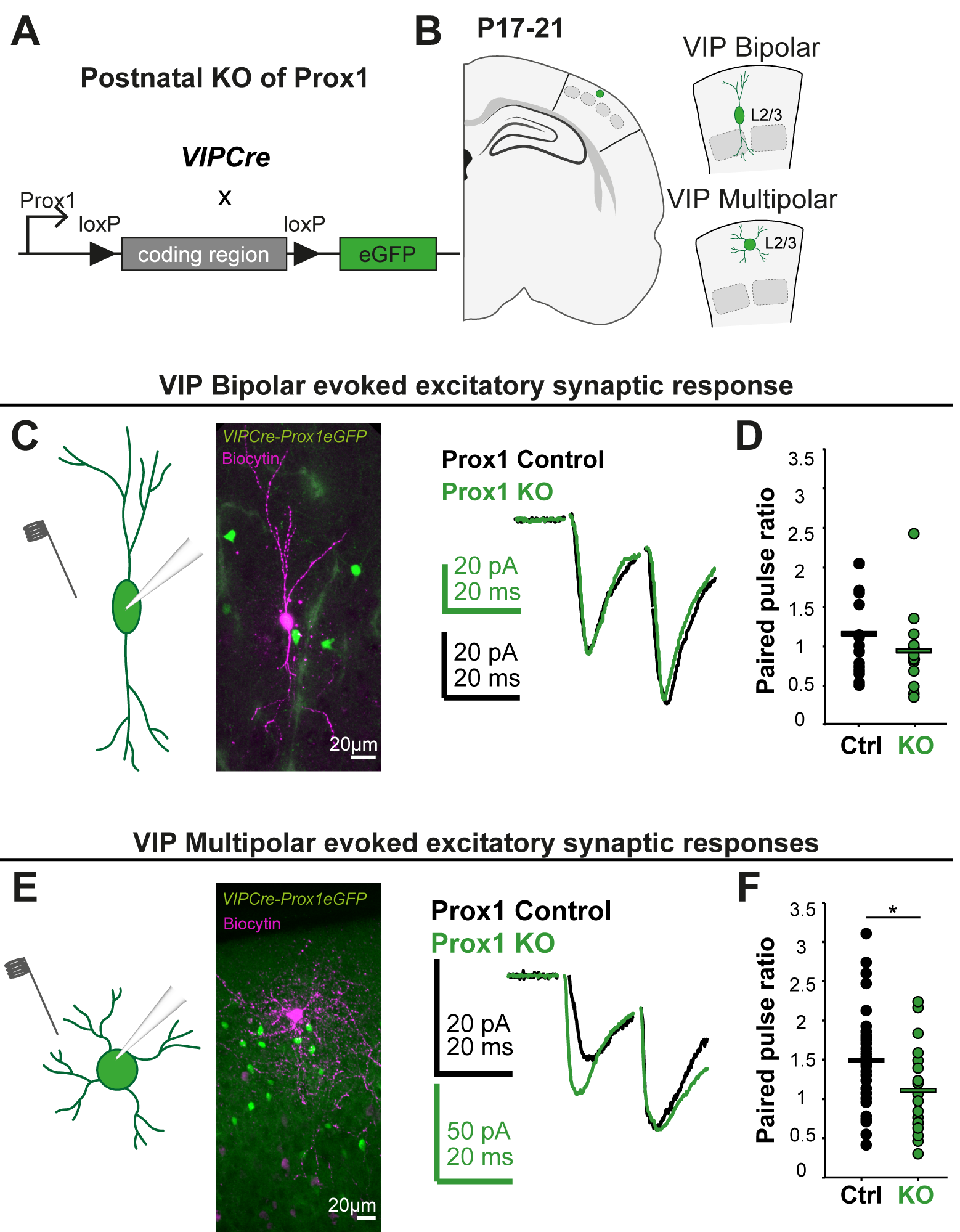
Postnatal Prox1 removal leads to changes in presynaptic release probability onto VIP multipolar but not bipolar cells. (A) Visual representation of Prox1 conditional knock-out strategy. VIPCre turns on postnatally and removes part of the coding region of the Prox1 locus shifting eGFP into frame and allowing for the visualization of the control (Prox1 het) and Prox1 KO cells. (B) Schematic representation of the experiment. L2/3 multipolar and bipolar VIP cells were recorded in the somatosensory barrel cortex at P17-21 of acute brain slices from control and Prox1 KO animals. (C) & (E) Left: schematic representation for probing the evoked excitatory synaptic responses onto control and Prox1 KO VIP cells. The stimulating electrode is shown close to the soma and proximal dendrites of the eGFP-positive cells, which are recorded in whole-cell patch clamp mode.Middle: the recorded cells were filled with biocytin and their morphology revealed post-hoc. Right: examples of a pair of evoked synaptic responses for control and KO cells, which are overlaid and scaled to the second response. (D) All data points for paired pulse ratio (PPR; 2^nd^/1^st^ response) for control (n/N=18/11) and KO (n/N=13/11) bipolar VIP cells, p = 0.3, t = 1.1. (F) All data points for paired pulse ratio of control (n/N=33/18) and KO (n/N=19/14) multipolar VIP cells, p = 0.03, t = 2.17. Statistics: t-test.

To reveal the mechanism by which Prox1 regulates the strength of excitatory inputs onto the two VIP subtypes we further examined the short-term dynamics of evoked synaptic responses onto control and Prox1 KO cells at P17-21 (Figure 1B). Changes in the paired pulse ratio (PPR) of the 2^nd^ to 1^st^ evoked EPSC amplitude implicate a presynaptic versus postsynaptic change, and provide an indication of whether presynaptic release probability for a given synapse is low (PPR > 1), moderate (PPR ∼ 1), or high (PPR < 1). Two electrical stimuli were delivered at 50 Hz through a glass pipette placed close to the soma and proximal dendrites of the recorded cells. We found that glutamatergic synapses onto control VIP bipolar cells show negligible synaptic facilitation (mean PPR: 1.15 ± 0.13) and that Prox1 KO does not affect the PPR significantly (mean PPR 0.93 ± 0.15) (Figure 1C, D). On the other hand, excitatory inputs onto VIP multipolar cells show more pronounced facilitation in the control condition (mean PPR: 1.51 ± 0.11) and a notable reduction of the PPR upon removal of Prox1 (mean PPR: 1.13 ± 0.1) (Figure 1E, F). These results demonstrate that in control conditions the initial release probability of glutamatergic synapses onto VIP multipolar cells is lower than onto bipolar cells, in line with previous reports showing that L2/3 CR+ bipolar cells display short-term synaptic depression at 10Hz and no change at 50Hz (Caputi et al., 2009) (https://portal.brain-map.org/explore/connectivity/synaptic-physiology). Importantly, the data also shows that postnatal removal of Prox1 leads to an increase in initial release probability of excitatory synapses onto multipolar, but not bipolar cells.

### 2) Loss of Prox1 leads to a downregulation of the trans-synaptic protein Elfn1

Having identified a Prox1-dependent suppression of synaptic release probability onto multipolar cells, we hypothesized that this may occur through the regulation of genes encoding for synaptic proteins. To identify such potential downstream targets of Prox1 we performed a RNA sequencing screen on control and Prox1 KO cells, after fluorescence activated cell sorting (FACS) of the GFP+ VIP cells at P8 and P12 (Figure 2A). Differential gene expression analysis of the data identified several potential candidate genes (Figure 2B, C) (Supplementary Figure 2C, D), which were analysed for gene ontology (GO) enrichment. The GO analysis revealed that most of the upregulated genes in Prox1 KO cells are associated with glial cell programs (Figure 2D). This result suggests that in control VIP cells, post-natal expression of Prox1 supresses those glial programs, directing the cell instead towards a neuronal fate. In line with this, as well as with our functional findings, the most enriched downregulated genes are associated with synapses and synapse-associated signalling (Figure 2D, E). Within those synapse-associated programs we specifically looked for genes that could mediate trans-synaptic interactions between the postsynaptic VIP and the presynaptic excitatory cell to regulate synaptic release. One such gene was the Extracellular Leucine Rich Repeat and Fibronectin Type III Domain Containing 1 (*Elfn1*) (Figure 2C), which creates the strongly facilitating excitatory synaptic inputs of SST INs (Sylwestrak and Ghosh, 2012). Postsynaptic Elfn1 produces cell-autonomous suppression of presynaptic glutamate release through its trans-synaptic recruitment of metabotropic-glutamate receptor 7 (mGluR7) (Stachniak et al., 2019; Tomioka et al., 2014).

**Figure 2:**
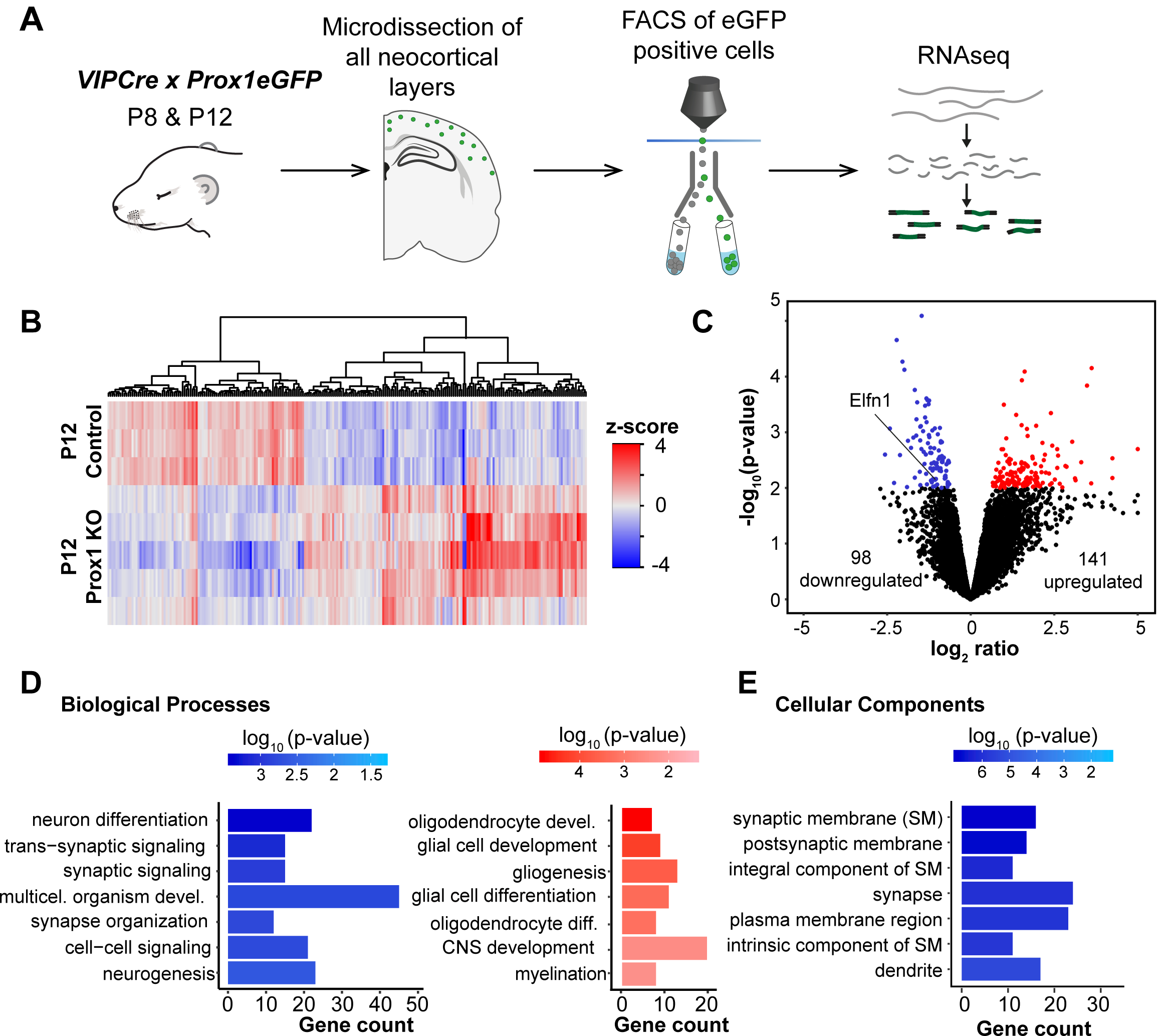
Postnatal removal of Prox1 from VIP cells leads to transcriptomic changes in synaptic proteins. (A) Schematic of experimental workflow. VIPCre-Prox1eGFP control and KO cells were sorted using FACS at P8 and P12 and bulk RNA sequencing was performed (B) Heat map showing the clustering according to function of up- (red) and downregulated (blue) genes at P12. (C) Volcano Plot highlighting the differentially expressed candidate genes at P12. They were selected based on log_2_ratio ≥|0.5| and p-value≤0.01 (D) GO: Biological Processes enrichment analysis of the candidate genes at P12. The down-regulated genes are depicted in blue and the up-regulated gene in red. (E) GO: Cellular Components (CC) enrichment analysis of the candidate genes at P12. The down-regulated genes are depicted in blue. The up-regulated genes did not show any clustering in the GO:CC

The RNA sequencing data showed that the level of *Elfn1* mRNA in VIP cells is high at both P8 and P12 (Supplementary Figure 3A). To assess if *Elfn1* is continuously expressed into adulthood, we consulted published single-cell RNA sequencing data (Tasic et al., 2018) and found that, in the adult cortex, almost all VIP cells express *Elfn1* (Supplementary Figure 3B). Furthermore, this dataset also shows a persistent expression of *Prox1*, which suggests a continued requirement for these two factors in the maintenance of cell function throughout life.

### 3) Reduction in Elfn1 expression recapitulates the Prox1 KO phenotype in VIP multipolar cells

To test if Elfn1 is the downstream molecule responsible for the synaptic phenotype in Prox1 KO multipolar VIP cells we used a compound mouse line that labeled VIP neurons (*VIPCre* x tdtomato reporter; *Ai14*) in the background of a germline *Elfn1* KO allele (*Elfn1KO*) (Figure 3A). We chose to compare VIP tdtomato+ cells from heterozygous *Elfn1KO* (Het Elfn1) animals to those from wildtype (control) littermates, to match our findings in the RNA sequencing screen, which showed a two-fold reduction of *Elfn1* in Prox1 KO cortex (Figure 2C). We found that reducing Elfn1 expression does not affect the PPR significantly in VIP bipolar cells (control mean PPR: 1.15 ± 0.14; Het Elfn1 mean PPR 0.98 ± 0.08) (Figure 3B, C). On the other hand, excitatory inputs onto VIP multipolar cells showed a notable reduction in the PPR when Elfn1 levels are reduced (control mean PPR: 1.30 ± 0.13; Het Elfn1 mean PPR: 0.88 ± 0.08) (Figure 3D, E). Our results show that a decrease in Elfn1 expression recapitulates the effect of knocking-out Prox1 in VIP multipolar cells but has no notable effect in bipolar cells. This finding suggests that Prox1 is important for initiating and/or maintaining the expression of Elfn1 in multipolar cells, which leads to facilitation of incoming excitatory synaptic responses.

**Figure 3:**
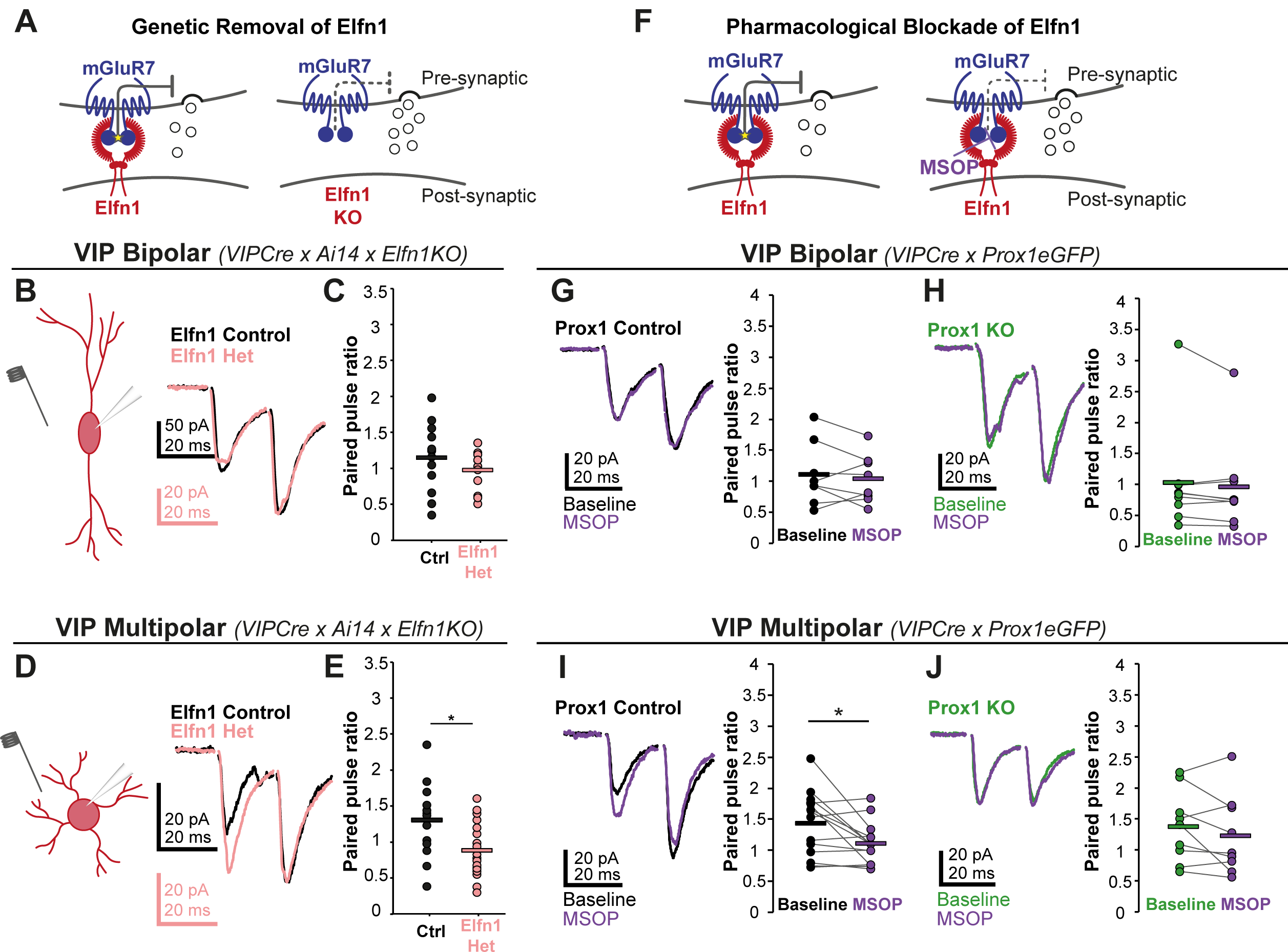
Elfn1 contributes to synaptic facilitation onto VIP+ multipolar, but not bipolar cells. (A) A compound mouse line labeling VIP cells was combined on the background of an *Elfn1* germline knock-out. The presence of Elfn1 increases initial release probablity of glutamate. Animals heterozygous (het) and wildtype (wt) for the germline removal were used as experimental and control group respectively. (B) /(D) Left: schematic representation of the evoked paired-pulse protocol, with the extracellular stimulating and intracellular recording electrodes shown. Right: example of a pair of synaptic responses evoked onto VIP bipolar and multipolar cells after stimulation (scaled to 2nd evoked response). (C) All data points for paired pulse ratio (PPR) of Elfn1 Control (n/N=12/4) and Het (n/N=12/4) bipolar VIP cells, p = 0.3, t = 1.06. (E) All data points for PPR of Elfn1 Control (n/N=14/6) and Het (n/N=20/8) multipolar VIP cells, p = 0.008, t = 2.84. Statistics: t-test (F) Testing for the effect of MSOP, a presynaptic mGluR blocker and hence Elfn1 function, on evoked excitatory responses onto control (Het) and Prox1 KO cells. (G-J) Left: example of a pair of synaptic responses evoked onto VIP bipolar and multipolar cells after stimulation (scaled to 2nd evoked response). Right: all data points for PPR are plotted. (G) PPR of bipolar Prox1 Control cells (n/N=8/8) under baseline conditions and with MSOP, p = 0.5, t = 0.74. (H) PPR of bipolar Prox1 KO cells (n(N=9/9) under baseline conditions and with MSOP, p = 0.2, t = 1.26. (I) PPR of multipolar Prox1 Control cells (n/N=14/9) under baseline conditions and with MSOP, p = 0.02, t = 2.74. (J) PPR of multipolar Prox1 KO cells (n/N=9/8) under baseline conditions and with MSOP, p = 0.2, t = 1.33. Statistics: paired t-test

### 4) Prox1-dependent engagement of Elfn1 in VIP cells

To directly test the relationship between Prox1 and the expression of Elfn1 in multipolar and bipolar cells, we turned to a pharmacological agent, (RS)-α-Methylserine-O-phosphate (MSOP), that acts as an antagonist for presynaptic mGluRs (including mGluR7), which are reported to interact trans-synaptically with Elfn1 (Figure 3F) (Dunn et al., 2018). Application of MSOP indirectly tests for the presence of Elfn1 in control and Prox1 KO VIP cells by blocking the constitutive suppression of synaptic release that Elfn1 induces through mGluRs (Stachniak et al., 2019). We therefore evoked excitatory synaptic events onto the two VIP subtypes, as described above, and assessed changes in neurotransmitter release in response to MSOP. We found that MSOP did not affect the release probability onto control (PPR: 1.11 ± 0.18 at baseline vs. 1.04 ± 0.14 in MSOP) or Prox1 KO VIP bipolar cells (PPR: 0.95 ± 0.29 at baseline vs. 0.88 ± 0.24 in MSOP) (Figure 3G, H). In contrast, MSOP markedly increased the initial release probability of excitatory inputs onto control VIP multipolar cells and thus reduced the paired pulse ratio (PPR: 1.43 ± 0.14 at baseline vs. 1.11 ± 0.09 in MSOP) (Figure 3I). However, the effect of MSOP was absent in the Prox1 KO VIP multipolar cells (PPR: 1.37 ± 0.19 at baseline vs. 1.23 ± 0.21 in MSOP) (Figure 3J).

In previous studies, both VIP bipolar and multipolar cells were shown to express *Elfn1* in the adult cortex (Paul et al., 2017; Tasic et al., 2018). This mRNA data stands in apparent contradiction to the VIP multipolar cell-selective effects of Elfn1 we observe in our Prox1 loss-of-function, Elfn1 downregulation, and pharmacological experiments. We therefore hypothesized that either only multipolar cells express *Elfn1* before P21, or that Prox1 selectively regulates *Elfn1* expression in multipolar cells. To test these two possibilities, we collected and sectioned brains from Prox1 conditional postnatal KO mice, (*Prox1fl* x *VIPCre* x *Ai14*) at P12 and performed *in situ* hybridization for *Elfn1* mRNA and *CR* mRNA to distinguish putative bipolar (*CR*+) and multipolar (*CR*-) VIP cells (Figure 4A, B). We then obtained images of whole cortices and analyzed them using a custom-written MATLAB script (see Methods). We first assessed the presence of *Elfn1* mRNA in control VIP cells at P12, which showed that, similarly to the adult cortex (Paul et al., 2017; Tasic et al., 2018), *CR*+ VIP cells express the gene at a higher level than *CR*-VIP cells (Figure 4C, Supplementary Figure 4C). This finding negates the first hypothesis regarding age-dependent selective expression of *Elfn1* in multipolar cells.

**Figure 4:**
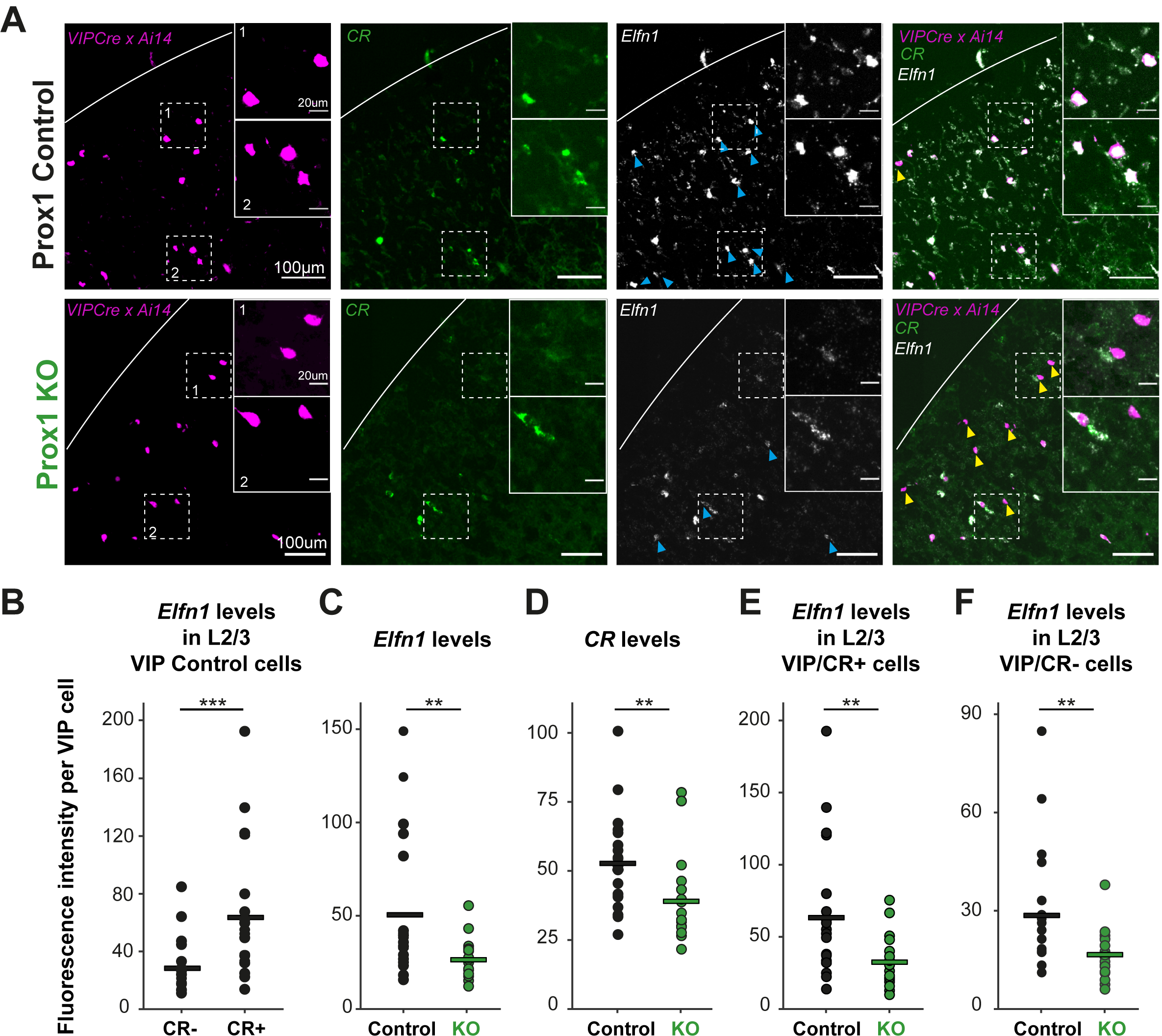
Prox1 regulats *Elfn1* expression in all VIP interneurons. (A) Examples of *Elfn1* and *CR in-situ* labelling in L2/3 tdtomato positive VIP cells in control (top) and Prox1 KO (bottom) tissue at P12. The insets show the close up of the dashed-line boxes including *CR* positive *(CR+)* and *CR* negative *(CR-)* VIP cells. The white parabolic line indicates the pia. Yellow arrows highlight VIP positive *Elfn1* negative cells. Blue arrows highlight *Elfn1* positive VIP cells. Note: there are also *Elfn1* positive cells that are not VIP positive likely corresponding to somatostatin cells. (B) *Elfn1* levels in VIP multipolar (*CR-*) and bipolar (*CR+*) cells in L2/3 of the somatosensory cortex, p=0.001 (N=3 animals, n=19 sections/images). (C) *Elfn1* levels in VIP cells in all layers and cortical areas in control (N/n=3/19) and Prox1 KO (N/n=2/18) tissue, p=0.004. (D) *CR* levels in VIP cells in all layers and cortical areas in control (N/n=3/19) and Prox1 KO tissue (N/n=2/18), p=0.004. (E) *Elfn1* levels in VIP bipolar cells in L2/3 of the somatosensory cortex in control (N/n=3/19) and Prox1 KO (N/n=2/18) tissue, p=0.004. (F) *Elfn1* levels in VIP multipolar cells in L2/3 of the somatosensory cortex in control (N/n=3/19) and Prox1 KO (N/n=2/18) tissue, p=0.006. Statistics: Mann-Whitney-U-Test.

We subsequently compared *Elfn1* and *CR* expression between control and KO tissue and found a clear reduction of both signals (Figure 4D, E), which is in line with the RNAseq results that showed a downregulation of both genes (log2 ratio of -1.466 for *CR* and -1.065 for *Elfn1*). Intriguingly, this reduction was seen for both *CR*+ (bipolar) VIP cells and *CR*-VIP ones (many of which belong to the multipolar subtype) (Figure 4F, G, Supplementary Figure 4D, E). These results suggest that Prox1 regulates *Elfn1* in both subtypes of VIP cells. Hence our second hypothesis is also negated, leaving open other possible mechanisms that are discussed below.

## Discussion

In the mammalian nervous system the TF Prox1 is known to regulate cell-cycle exit (Cid et al., 2010; Kaltezioti et al., 2010) and cell fate determination (Iwano et al., 2012; Kaltezioti et al., 2014) of neural precursor cells. Post-mitotic removal of Prox1 in embryonic CGE-derived cortical INs results in a failure of CR+ VIP INs to migrate to the correct cortical layers, a dramatic decrease in their numbers, and a subsequent reduction in the excitatory synaptic input onto the remaining cells (Miyoshi et al., 2015). Postnatal removal of Prox1 circumvents cell death and layer mis-targeting. Nevertheless, we find a continued requirement for the TF in the regulation of the proper synaptic integration and final specification of VIP subtypes. Our results demonstrate that post-natal expression of Prox1 is necessary for maintenance of expression levels of the bipolar cell marker *CR*, and for both VIP bipolar and multipolar cells to connect properly to their cortical networks. Furthermore, we show that Prox1 is necessary for synaptic facilitation of excitatory inputs onto VIP multipolar cells and that it exerts this function by regulating Elfn1 expression. An emerging theme from single cell transcriptomics studies is that cell adhesion molecules like Elfn1 may determine the cell specific interactions that guide connectivity profiles of IN subtypes (Paul et al., 2017). Accordingly, by regulating VIP subtype specific Elfn1 engagement, the TF Prox1 promotes and maintains functional diversity for VIP IN subtypes.

The selective engagement of Elfn1 in VIP multipolar cells leads to excitatory synaptic facilitation through a reduction in the initial release probability. In SST INs this Elfn1-dependent facilitation of excitatory inputs prevents rapid recruitment, thereby creating the characteristic “delayed” firing of these INs (Sylwestrak and Ghosh, 2012) and biasing responsiveness towards high frequency activity (Pouille and Scanziani, 2004; Stachniak et al., 2019). The same mechanism would allow VIP multipolar cells to selectively tune to the high frequency activity characteristic of cortico-cortical communication (Palmer et al., 2012). Thus, by regulating Elfn1, Prox1 may prime these neurons to coordinate intra-cortical communication within the superficial layers of the cortex. Interestingly, the SST INs do not express Prox1, therefore our results not only identify a novel functional role for Elfn1 in VIP multipolar INs, but also reveal a novel regulatory pathway for Elfn1 expression.

In contrast, short-term plasticity of excitatory inputs onto bipolar cells is unaffected by removal of Prox1, by decreased Elfn1 expression, or by the pharmacological blockade of its synaptic effects. These findings are surprising given that our results and published data show high levels of *Elfn1* mRNA expression in bipolar VIP cells. Thus, it appears that in this VIP subtype, *Elfn1* mRNA levels are uncoupled from Elfn1 function. An explanation for this discrepancy between expression and function could be that the presynaptic excitatory terminals onto bipolar cells lack mGluR7, despite the presence of Elfn1 protein. This could arise if distinct cell populations target VIP bipolar and multipolar cells, which, for example, may receive different amounts of thalamo-cortical and cortico-cortical inputs. The two subtypes tend to sit in different parts of L2/3, with VIP multipolar cells often found close to the border to L1, while VIP bipolar cells usually are located closer to L4 (He et al., 2016). In combination with their distinct dendritic architecture, this could determine the origin of inputs the VIP subtypes receive (Sohn et al., 2016). Additionally, the distinct glutamate receptor composition expressed by the two VIP subtypes (Paul et al., 2017) could also point to cell selective synaptic wiring. Alternatively, the cell type specificity we observe could be a combination of Prox1-dependent and independent mechanisms that would regulate subtype specific protein expression, such as differential microRNA expression or alternative splicing. Indeed, *Elfn1* can be regulated by microRNAs, and showed a 75% increase in expression following deletion of the *mirg* microRNA cluster in an induced neuronal culture system (Whipple et al., 2020).

Even though we find that Prox1 does not affect release probability onto bipolar cells, it clearly plays a role in the cells’ synaptic integration into the circuit. Previous work demonstrated a critical role for electrical activity in the final specification of VIP bipolar cells (De Marco García et al., 2011). This finding may well relate to the differential expression of excitatory synaptic components such as the subunits of NMDA receptors. Published research has shown that adult cortical bipolar VIP cells have high expression of the NR2B subunit (Paul et al., 2017). This finding is intriguing given that the receptors containing this subunit are critical for the integration of superficial Reelin-positive INs, via their activation by thalamocortical terminals (De Marco García et al., 2015). Although we find no evidence for direct regulation of NR2B expression by Prox1, the regulation of cofactors that control thalamocortical excitation through NR2B-containing receptors could underlie the observed Prox1 KO phenotype in bipolar cells.

The final specification of neurons is a process that takes place after the cells’ birth, as they start integrating into the resident circuit. It is during this time of establishing connections that the needs of the circuit, as defined by those of the animal, can instruct a neuron towards its mature state, which includes the proper construction and function of inputs and outputs. In inhibitory INs these input/output specificities vary considerably, not only between cardinal IN classes, but also between the subtypes within a class (Huang and Paul, 2019). Our data provides evidence that the continuous expression of the TF Prox1 is necessary for the final specification of bipolar and multipolar VIP cells, allowing them to acquire diverging roles within the adult cortical network.

## Material and Methods

### Mice

All animal experiments were approved by the Cantonal Veterinary Office Zurich and followed Swiss national regulations. Animal lines used in this study are: *VIPCre* (Vip^tm1(cre)Zjh/J^) (Taniguchi et al., 2011), *Ai14* (B6;129S6-Gt(ROSA)^26Sortm14(CAG-tdTomato)Hze/J^) (Madisen et al., 2010), *Prox1fl* (Prox1^tm2Gco^) (Harvey et al., 2005), *Prox1eGFP* (Prox1^tm1.1Fuma^) (Iwano et al., 2012) and *Elfn1KO* (Elfn1^tm1(KOMP)Vlcg^) (created by the Knock Out Mouse Project). The following compound lines were created for this study: *VIPCre x Prox1eGFP* and *VIPCre x Ai14 x Elfn1KO* for electrophysiology experiments, *VIPCre x Ai14 x Prox1fl* for *in situ* experiments. All crosses were setup to produce both KO and Control animals in the same litter and littermate controls were used throughout the study.

### Electrophysiology

Whole-cell patch-clamp electrophysiological recordings were performed on GFP labelled VIP neurons located in neocortical layers II–III of barrel cortex (approximately bregma -0.5 to -2.0 mm) in acute brain slices prepared from P17–P22 male and female mice. Briefly, animals were decapitated and the brain was dissected out and transferred to cold cutting solution containing (in mM): 75 sucrose, 87 NaCl, 25 NaHCO_3_, 25 D-glucose, 2.5 KCl, 7 MgCl_2_, and 1.25 NaH_2_PO_4_, aerated with 95% O_2_ / 5% CO_2_. 300 μm slices were recovered in artificial cerebrospinal fluid (ACSF) composed of (in mM): 128 NaCl, 3 KCl, 26 NaHCO_3_, 1.25 NaH_2_PO_4_, 1 MgCl_2_, 2 CaCl_2_, and 10 glucose at 34°C for 15 min. Acute slices were perfused at a rate of 2–3 ml/min with oxygenated recording ACSF at room temperature. Patch electrodes were made from borosilicate glass (Harvard Apparatus) and had a resistance of 2-4 MΩ. The intracellular solution contained (in mM): 126 cesium methanesulfonate, 4 CsCl, 10 HEPES, 20 phosphocreatine, 4 MgATP, 0.3 NaGTP, pH 7.3, 290 mOsm, with addition of 2.5 mg/mL of biocytin.

Experiments were performed in voltage-clamp mode using the Axopatch 200B amplifier (Molecular Devices). sEPSCs were recorded at a holding potential (Vh) = -65 mV, with a sampling rate of 10 kHz and were filtered on-line at 2 kHz. Access resistance was monitored to ensure the stability of recording conditions. Recordings with access resistance >40 MΩ, whole cell capacitance <4 pF or holding current >200 pA were excluded. No compensation was made for access resistance and no correction was made for the junction potential between the pipette and the ACSF. Following a baseline stabilization period (3 min), synaptic currents recorded in 2x 3 min traces were low pass filtered at 400 Hz, then analyzed using clampfit event detection template match. Evoked synaptic responses were recorded at Vh= -70 mV. Electrical stimulation from a Digitimer isolated stimulator (DS2A Mk.II) was through a monopolar glass pipette (2-4 MΩ) positioned in L2/3. The stimulating electrode was placed typically 50 - 150 μm from the recorded cell, parallel to the pial surface. Stimulation intensity and duration were adjusted to produce stable evoked EPSC amplitudes. Stimulation intensities were larger for bipolar cells (86 ± 1V for 103 ± 6 µs) than for multipolar cells (74 ± 2V for 86 ± 6 µs, p= 3 x 10^−6^, p= 0.06), but did not differ between genotypes (multipolar prox1 control: 69 ± 3V for 57 ± 5 µs, KO: 71 ± 4V for 74 ± 10 µs, p= 0.6, p= 0.2; bipolar prox1 control: 85 ± 2V for 91 ± 9 µs, KO: 84 ± 3V for 88 ± 15 µs, p= 0.9, p= 0.9; multipolar Elfn1 control: 77 ± 3V for 114 ± 13 µs, Het: 82 ± 3V for 125 ± 19 µs, p= 0.2, p= 0.6; bipolar Elfn1 control: 86 ± 3V for 113 ± 13 µs, Het: 89 ± 2V for 130 ± 15 µs, p= 0.3, p= 0.4, t test). Paired pulse ratios (PPR) were calculated from an average of 12 sweeps, after a 2-minute stable baseline was established. Bath applied compound MSOP (100 µM, pH 7.4) was dissolved in water.

### Electrophysiology data analysis

Values are represented as mean ± SEM. Number of measurements n/N indicates cells recorded (n) from animals (N), typically using 1 cell per slice to recover biocytin-stained cell morphology. Cell type was classified as bipolar vs. multipolar based on cell body morphology and number of dendritic processes emanating from it, using both initial determination prior to patching and post-hoc verification of recovered biocytin-labelled cells. Matched recordings were performed with Prox1 control and Prox1 KO littermates on the same day, whenever possible. Statistical testing was done in Matlab. Comparisons within conditions were made by two-tailed paired Student’s t-test, treatment versus baseline. Comparisons across conditions or between genotypes were done with an unpaired t-test assuming unequal variance. To accommodate skewed distributions in spontaneous synaptic frequency and amplitude, we performed Kolmogorov-Smirnoff non-parametric testing on event distributions and Wilcoxon non-parametric testing on averages (mean frequency, median amplitude). For multiple comparisons, a one-way or two-way ANOVA was done with a Bonferroni post hoc test. Statistical outcomes are represented in figures as: n.s. p > 0.05, *p < 0.05, **p < 0.01, ***p < 0.001.

### Cell sorting and RNA sequencing

EGFP-labeled cells were purified from the P8 and P12 cortices of control heterozygous and Prox1 KO animals *(VIPCre x Prox1eGFP/+* and *VIPCre x Prox1eGFP/eGFP*) as described (Hempel et al., 2007). Briefly, animals were anesthetized in 2% vol/vol isoflurane (∼1.5min), decapitated, the brain was extracted, the olfactory bulb and cerebellum were removed, and the brain was cut into 400-um thick coronal sections using a vibratome (Leica VT1000S; Leica, speed 5, vibration frequency 7-8), while in bubbling ice cold ACSF (recipe as above, with 1mM CaCl2 and 1mM MgCl2) in the chamber. Sections were collected and transferred into a protease digestion solution of ACSF (with Pronase (1mg ml-1), for 25 minutes and then transferred into a quenching solution of ACSF (with 1%FBS). Microdissection of the cortex to include the somatosensory areas was performed using a fluorescent dissection microscope (Olympus, MVX 10). Dissected cortices were collected in a 15mL falcon tube containing 1.5mL of ACSF sorting solution containing 1%FBS and DNase and cells were dissociated by gently triturating ten times with a large, then medium, and finally small fire-polished Pasteur pipette while avoiding the generation of bubbles. The cell suspension was then passed through a 50uM filter (Sysmex CellTrics) before automated fluorescence-activated cell sorting (FACS) using the MoFlo™ or FACSAria™ devices. A GFP negative littermate control cortex was also included as a negative control for the FACS setup. Cells were collected into Arcturus Picopure extraction buffer and immediately processed for RNA isolation using the Arcturus PicoPure Isolation Kit (KIT0204). RNA quality and quantity was measured using Agilent High Sensitivity RNA ScreenTape system (High Sensitivity RNA Screen Tape 5067-5579-5580-5581). All samples had high quality scores between 6-8 RIN. 4-5 RNA samples of each genotype for each of the two age groups (P8 and P12) were used to prepare 19 barcoded libraries.

The libraries were prepared using the Smart-seq2 protocol (Picelli et al., 2013). Briefly, total RNA was placed in 4 µl of lysis buffer (0.1% vol/vol Triton X-100, 2.5 mM dNTPs, 2.5 µM oligo-dT, 1 U/µl Promega RNasin Plus RNase inhibitor). Reverse transcription was performed followed by cDNA amplification. The quality of the cDNAs was evaluated using an Agilent 2100 Bioanalyzer. The resulting cDNA (1 ng) was fragmented using Illumina Nextera XT according to standard protocol. Nextera adapters containing Unique Dual Indices (UDI) were added by PCR. The libraries were double-sided size, selected and quantified using an Agilent 4200 TapeStation System.

TruSeq SR Cluster Kit HS4000 (Illumina, Inc, California, USA) was used for cluster generation using 10 pM of pooled normalized libraries on the cBOT. Sequencing was performed on the Illumina HiSeq 4000 single end 100 bp, using the TruSeq SBS Kit HS4000 (Illumina, Inc, California, USA).

### RNA sequencing data analysis

The raw reads were first cleaned by removing adapter sequences, trimming low quality ends, and filtering reads with low quality (phred quality <20) using Trimmomatic (Version 0.33) (Bolger et al., 2014). The read alignment was done with STAR (v2.5.3a)(Dobin et al., 2013) As reference we used the Ensembl genome build GRCm38. with the gene annotations downloaded on 2015-06-25 from Ensembl. The STAR alignment options were “--outFilterType BySJout --outFilterMatchNmin 30 --outFilterMismatchNmax 10 --outFilterMismatchNoverLmax 0.05 --alignSJDBoverhangMin 1 --alignSJoverhangMin 8 --alignIntronMax 1000000 --alignMatesGapMax 1000000 --outFilterMultimapNmax 50”.

Gene expression values were computed with the function featureCounts from the R package Rsubread (v1.26.0) (Liao et al., 2013). The options for featureCounts were: min mapping quality 10 - min feature overlap 10bp - count multi-mapping reads - count only primary alignments - count reads also if they overlap multiple genes. One sample was excluded from further analysis based on quality control standards (Supplementary Figure 2A, B).

To detect differentially expressed genes we applied a count based negative binomial model implemented in the software package EdgeR (R version: 3.6.0, EdgeR version: 3.26.1) (Robinson et al., 2009). The differential expression was assessed using an exact test adapted for over-dispersed data. Genes showing altered expression with adjusted (Benjamini and Hochberg method) p-value <0.05 were considered differentially expressed.

A list of potential candidate genes at both P8 and P12 was generated by selecting genes that had a log2ratio≥|0.5| and p-value≤0.01. This list was analysed for gene ontology (GO) enrichment using g:Profiler (Raudvere et al., 2019) (https://biit.cs.ut.ee/gprofiler/gost).

### *In situ* experiments

*VIPCre* x *Ai14* x *Prox1fl* animals were sacrificed at P12. In short, the animals were deeply anesthetized before transcardial perfusion with 1x PBS followed by ice-cold 4% PFA. The brains were dissected and post-fixed in ice-cold 4% PFA for 1h before being placed in a 30% Sucrose solution at 4°C for >24h for cryo-protection. The brains were embedded in OCT using a peel-away mold and stored at -80°C. Coronal 20µm-think brain sections containing barrel cortex were cut and collected on-slide using a cryostat (Microm International, HM 560M) and the slides were stored at -80°C until further processing.

*In situ* hybridization for *Elfn1* mRNA and *CR* mRNA was performed using the RNAscope kit (RNAscope Intro Pack for Multiplex Fluorescent Reagent Kit v2-Mm, 323136). Briefly, the slides were thawed, OCT residue was removed using 1x PBS (3×5min washes). The slides were then baked for 30mins at 60°C, post-fixed for 30mins in 4% PFA, dehydrated in an Ethanol dilution series (50%, 70%, 2x 100%) and incubated with RNAscope Hydrogen Peroxide for 10mins. RNAscope 1x Target Retrieval Reagents was brought to the boil and the slides were submerged for 2mins. Protease III was added to the slides and left to incubate for 45mins at 40°C. After the pre-treatment the probes (Mm-Calb2, 313641 and Mn-Elfn1, 449661-C3) were hybridized to the slices for 2h at 40°C and the signal was amplified using branched DNA amplification methods and visualized with Opal dyes (Opal 520 FP1487001KT and Opal690 FP1497001KT).

Following the RNAscope protocol the slices were immunostained to retrieve the tdtomato signal that was lost during the RNAscope protocol. Slides were washed two times with 1x PBS for 5min before being blocked for 30mins using 10% normal donkey serum and 1% bovine serum albumin (BSA) in 1xPBS. The primary antibody against tdtomato (goat anti tdTomato, SICGEN Antibodies, AB8181-200) was diluted 1:700 in 1xPBS-1%BSA and left to incubate at 4°C overnight followed by a 2h incubation of secondary antibody at room temperature (donkey anti goat Cy3, Jackson Immuno Research 705-165-147, diluted 1:1000 in 1xPBS-1%BSA). Slides were cover-slipped with Fluoromount-G with DAPI (00-4959-52) and imaged using a Slidescanner (Zeiss, AxioScan Z1). Mosaic images of the whole cortex were taken using a 20x objective.

### *In situ* data analysis

Image analysis was performed using custom written MATLAB codes. A two-dimensional difference of Gaussian feature enhancement algorithm was used to improve the VIP-tdtomato images, followed by Otsu thresholding to get an initial segmentation of neuronal cell bodies. To ensure an accurate representation of the cell body, segmentation of VIP INs was finalized using active contour segmentation (Kass et al., 1988) (MATLAB functions fspecial, conv2, graythresh and activecontour were used for segmentation).

During preprocessing of the *CR* and *Elfn1* images background subtraction was performed using a disk-shaped structuring element (MATLAB functions fspecial and imfilter were used for background subtraction). Subsequently, fluorescent intensity levels of both *CR* and *Elfn1* were measure within each of the segmented VIP IN cell bodies. Layers 1-6, as well as the somatosensory region, were segmented manually using the DAPI channel as reference (Supplementary Figure 4A, B) (MATLAB functions bwlabel, bwconncomp, regionprops were used for manual segmentation). To define whether a VIP cell is *CR*+ we followed a Bayesian approach by assuming 80% of VIP cells are *CR*+ (Kubota et al., 2011). Hence, once mean CR intensity per cell was calculated for each image, the population was thresholded such that approximately 80% of total VIP cells are considered CR positive. Statistical analysis of the data was done using the Mann-Whitney U test.

### Data and Code availability

Data and custom written codes are available upon request.

## Author Contribution

Conceptualization & Methodology T.J.S., R.K. and T.K.; Formal Analysis T.J.S., R.K. and A.Ö.A.; Investigation T.J.S., R.K. and O.H.; Writing-Original Draft, R.K.; Writing-Review & Editing, T.J.S., R.K., O.H., A.Ö.A. and T.K.; Supervision & Funding Acquisition, T.K.

## Acknowledgments

We would like to thank G. Fishell for helpful discussions and for supporting the initiation of this project in his lab, while at NYU Medical Center, by providing access to the Prox1fl and Prox1eGFP mice. We would also like to thank L. Ibrahim for her comments and efforts on this project. Furthermore, we thank F. Matsuzaki for his generous gift of the conditional Prox1eGFP mouse line (http://www2.clst.riken.jp/arg/mutant%20mice%20list.html CDB0482K). We are grateful to C. Aquino and L. Opitz at the Functional Genomics Center Zurich (FGCZ) who performed the sequencing of the extracted mRNA and data analysis. Slidescanner imaging was performed with equipment maintained by the Center for Microscopy and Image Analysis (ZMB), University of Zurich. This work was supported by grants from the European Research Council (ERC, 679175, T.K), the Swiss National Science Foundation (SNSF, 31003A_170037, T.K), Fond zur Förderung des Akademischen Nachwuchs of the UZH Alumni (T.K) and the Swiss Foundation for Excellence in Biomedical Research (R.K and T.K). O.H was supported by an EMBO post-doctoral fellowship.

## Additional Information

Supplementary Figures S1-S4

The authors declare no competing interests

