## Supplementary Figures for "Post-mitotic Prox1 expression controls the final specification of cortical VIP interneuron subtypes"

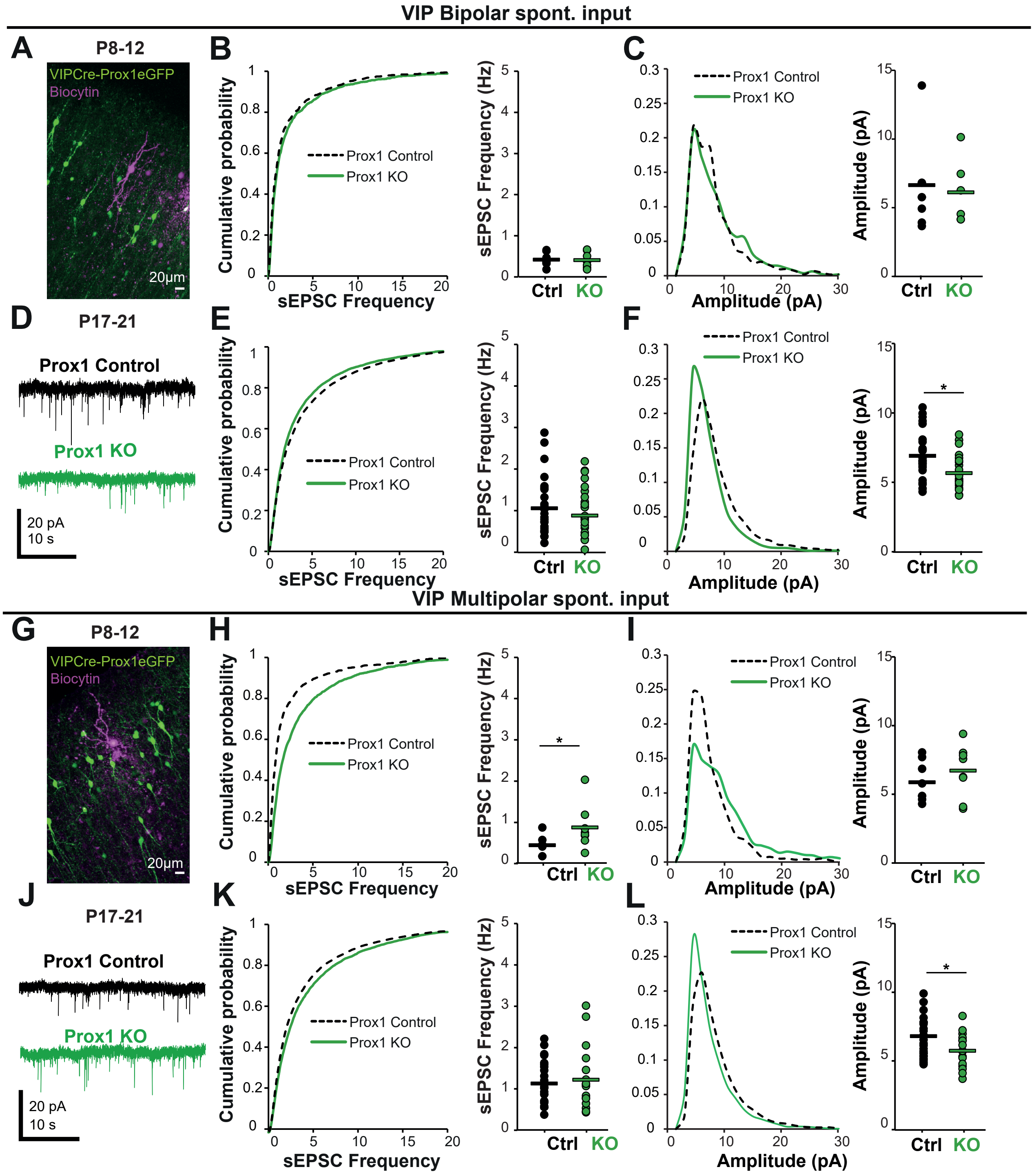


**Supplementary Figure 1: Prox1 differentially regulates synaptic connectivity onto VIP bipolar and multipolar cells**

(A) & (G) Example of patched control and KO cells at P8-12, shortly after Prox1 removal by Cre (~P3).

(B) & (C) Normalized distribution and mean/ median of sEPSC Frequency (B) and Amplitude (C) at P8-12 in control (n/N=7/6) and KO (n/N=7/6) bipolar VIP cells. Bipolar neurons showed no change in sEPSC frequency at this early time point. Frequency distribution: p=0.1, mean frequency: p=1, amplitude distribution: p=0.01, median amplitude: p=1.

(D)&(J) Example traces of control and KO cells at P17-21.

(E) & (F) Normalized distribution and mean / median of sEPSC Frequency (E) and Amplitude (F) at P17-21 in control (n/N=25/16) and KO (n/N=24/17) bipolar VIP cells. A leftward shift in sEPSC frequency distribution and amplitude distribution suggest reductions in excitatory synapse number and size. Frequency distribution: p = 1 x 10-9, mean frequency: p=0.8, amplitude distribution: p = 2 x 10-132, median amplitude: p=0.007.

(H) Normalized distribution and mean/ median of sEPSC Frequency (H) and Amplitude (I) at P8-12 in control (n/N=8/5) and KO (n/N=8/5) multipolar cells. Multipolar VIP neurons show an enhancement of synaptic excitation. A rightward shift in synaptic frequency distribution and a relative increase in large synaptic amplitudes suggest an enhancement of release probability. Frequency distribution: p = 4 x 10-38, mean frequency: p = 0.04, amplitude distribution: p = 4 x 10-31, median amplitude: p = 0.6.

(K) & (L) Normalized distribution and mean/ median of sEPSC Frequency (H) and Amplitude (I) at P17-21 in control (n/N=25/14) and KO (n/N=17/12) multipolar cells. P21 multipolar VIP neurons show a mixed phenotype: a rightward shift in synaptic frequency distribution as for P12, but with a leftward shift in amplitude distribution and reduction in median amplitude similar to P21 bipolar neurons. Frequency distribution: p = 3 x 10-12, mean frequency: p = 1, amplitude distribution: p = 2 x 10-27, median amplitude: p = 0.046.

Mean and Median statistics: Wilcoxon Rank Sum test, Distribution statistics: Kolmogorov-Smirnov test

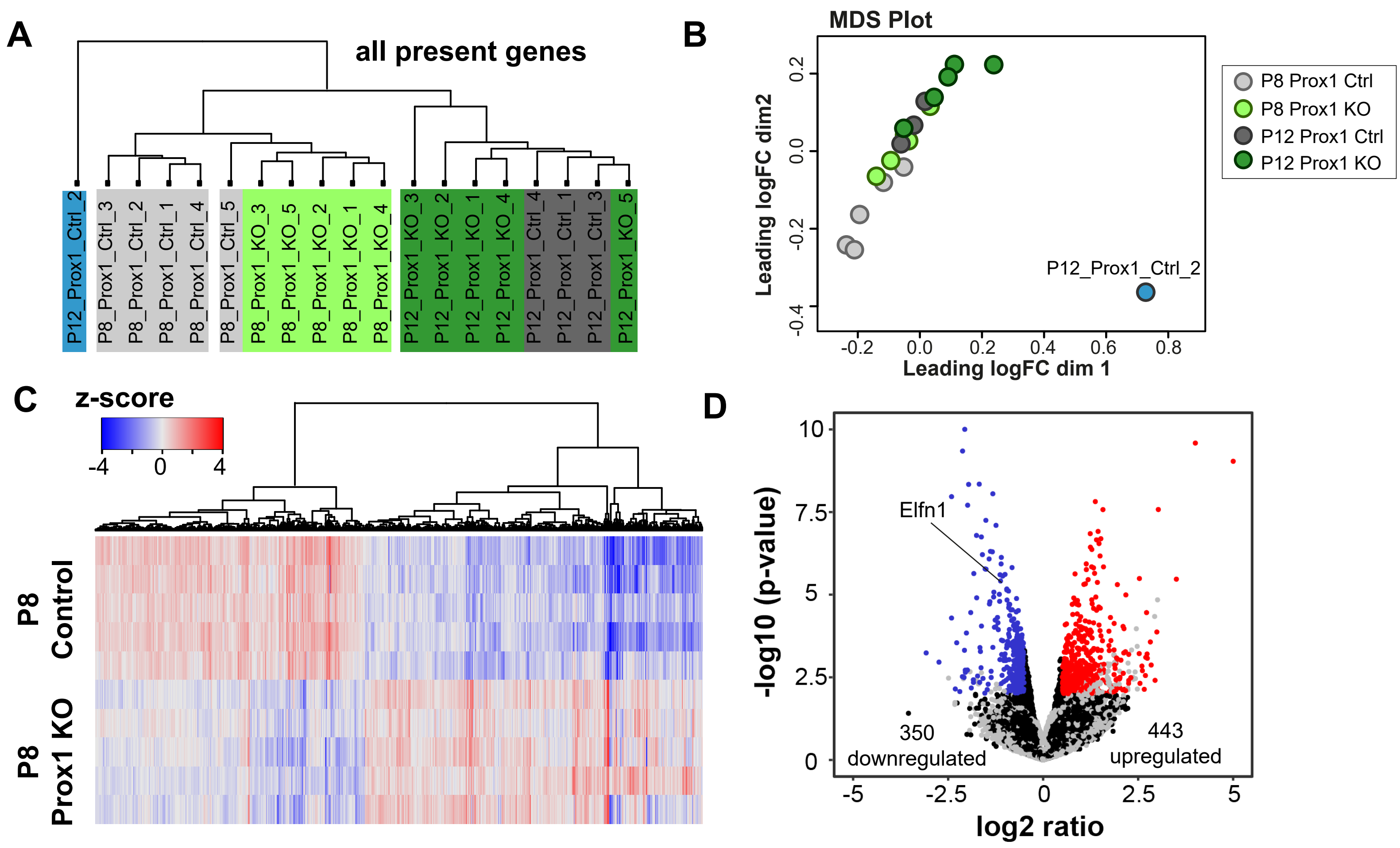


**Supplementary Figure 2: Validation of RNAseq Experiments & differential expression at P8**

(A) Clustering of all sequenced samples

(B) Multidimensional scaling (MDS) plot of all the sequenced samples.

(C) Heat map showing the clustering according to function of up- (red) and downregulated (blue) genes at P8

(D) Volcano Plot highlighting the candidate genes at P8. They were selected based on log2ratio≥|0.5| and p-value≤0.01

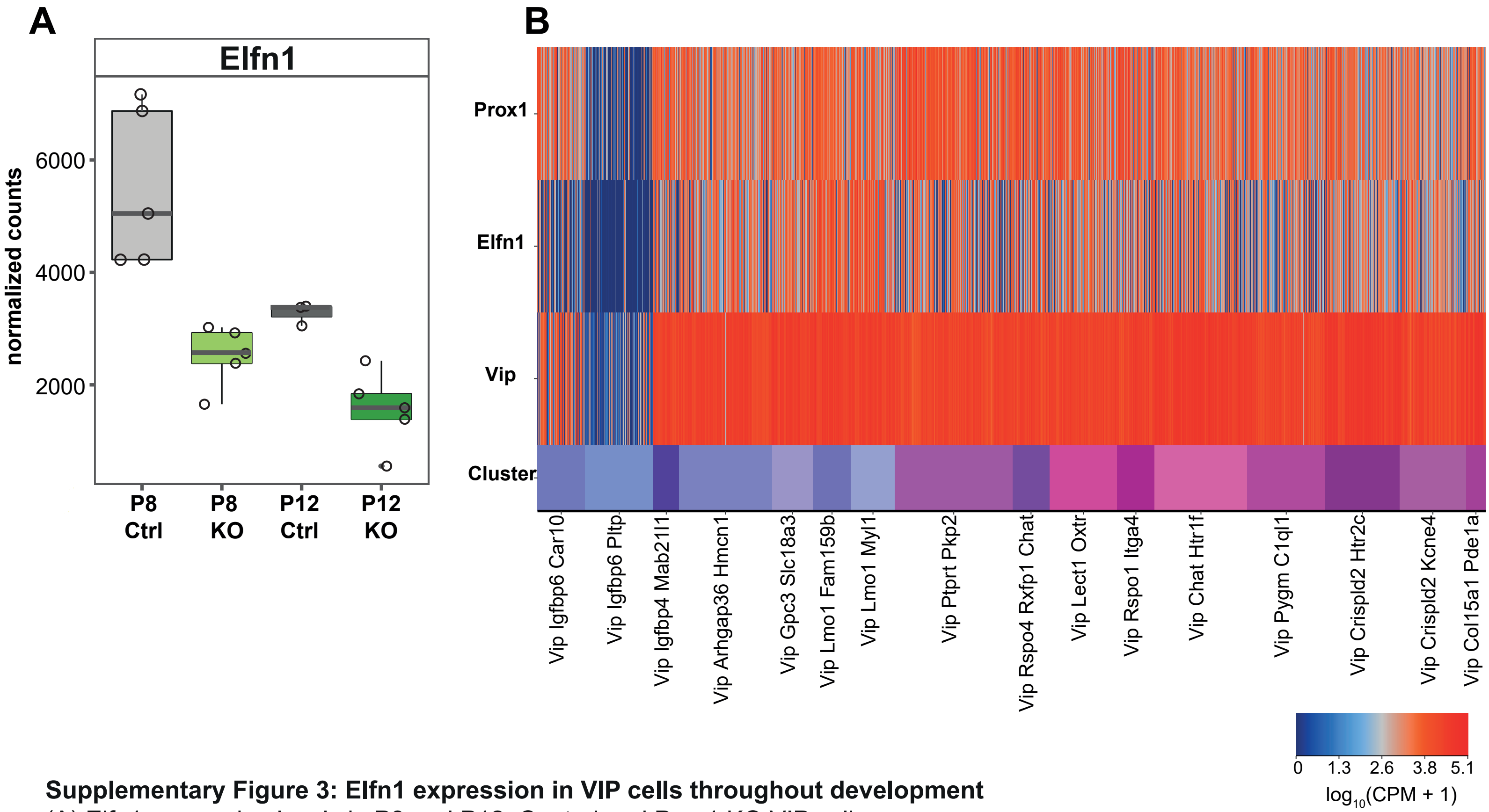


**Supplementary Figure 3: Elfn1 expression in VIP cells throughout development**
(A) Elfn1 expression levels in P8 and P12 Control and Prox1 KO VIP cells.
(B) Elfn1 and Prox1 expression levels in P50+ VIP cells. Single cell sequencing data from Tasic et al (2018).

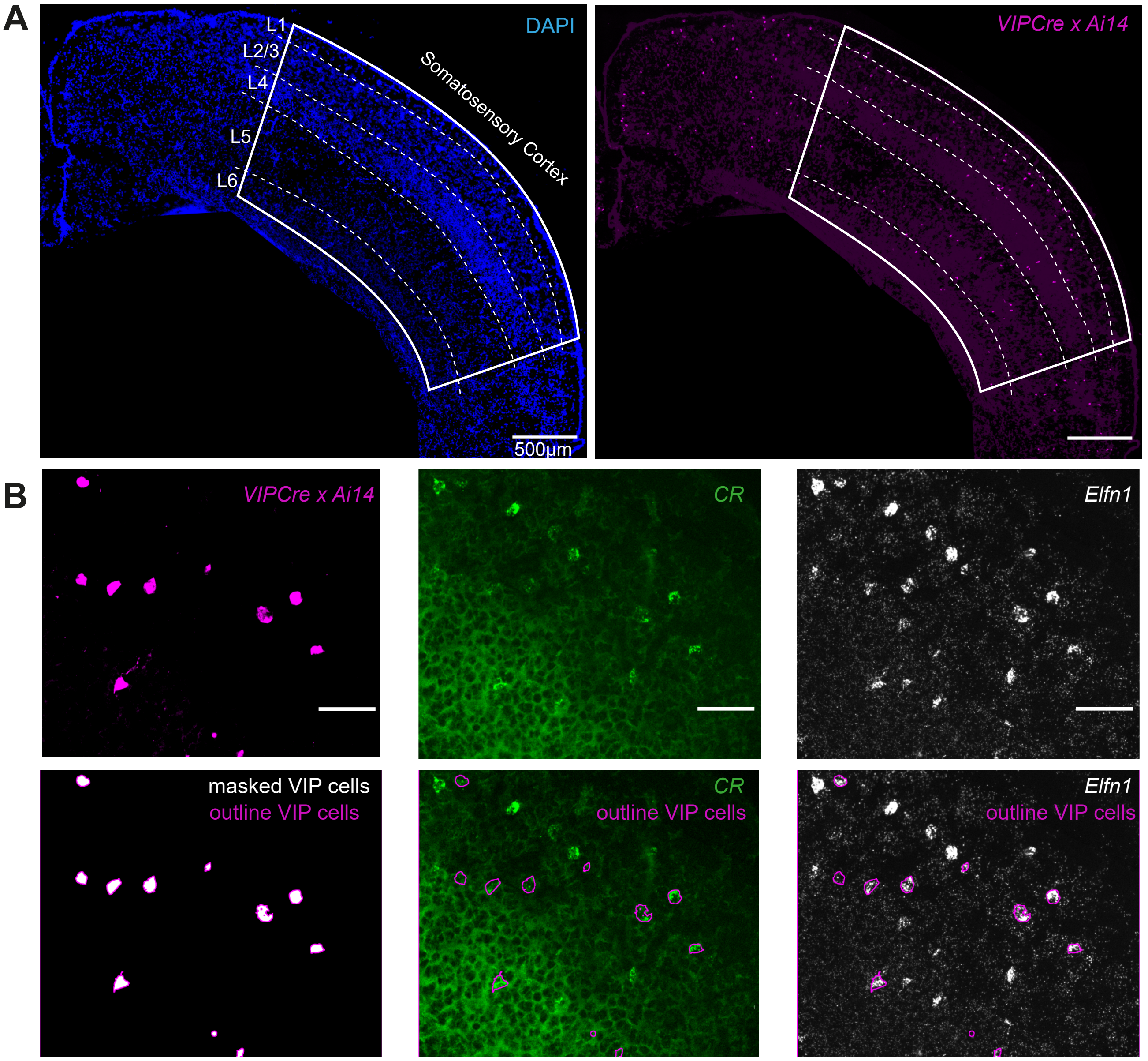


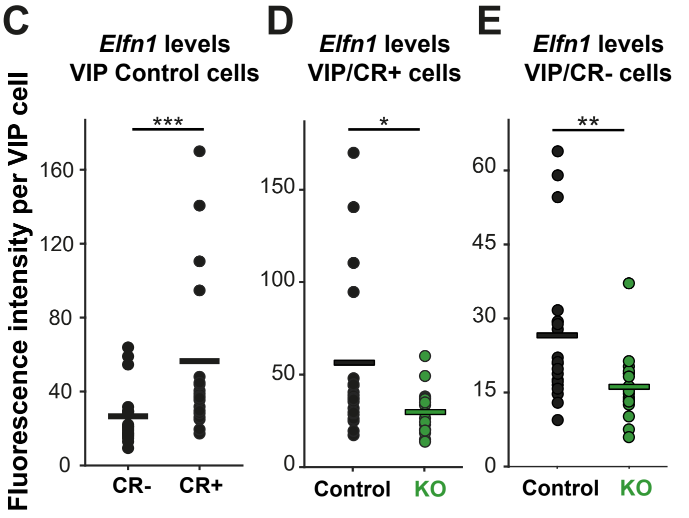


**Supplementary Figure 4: Elfn1 expression in VIP sutypes**

(A) Segmentation of somatosensory cortex and layers 1-6 based on the DAPI channel.
(B) Schematic representation of the analysis method: VIP cells were segmented and *CR* and *Elfn1* fluorescence levels were measured within the boundiers of the cell.
(C) *Elfn1* levels in VIP multipolar (*CR*-) and bipolar cells (*CR*+) in all layers of the somatosensory cortex at P12, p=0.001, (N=3 animals, n=19 slices/images).
(D) *Elfn1* levels in VIP bipolar cells in all layers of the somatosensory cortex in control (N/n=3/19) and Prox1 KO (N/n=2/18) tissue at P12, p=0.002.
(E) *Elfn1* levels in VIP multipolar cells in all layers of the somatosensory cortex in control (N/n=3/19) and Prox1 KO (N/n=2/18) tissue at P12, p=0.006.
Statistics: Mann-Whitney-U-Test.
